# Electrostatic regulation of the *cis*- and *trans*-membrane interactions of synaptotagmin-1

**DOI:** 10.1101/2022.11.29.518389

**Authors:** Houda Yasmine Ali Moussa, Yongsoo Park

## Abstract

Synaptotagmin-1 is a vesicular protein and Ca^2+^ sensor for Ca^2+^-dependent exocytosis. Ca^2+^ induces synaptotagmin-1 binding to its own vesicle membrane, called the *cis-*interaction, thus preventing the *trans*-interaction of synaptotagmin-1 to the plasma membrane. However, the electrostatic regulation of the *cis*- and *trans*-membrane interaction of synaptotagmin-1 was poorly understood in different Ca^2+^-buffering conditions. Here we provide an assay to monitor the *cis*- and *trans*-membrane interactions of synaptotagmin-1 by using native purified vesicles and the plasma membrane-mimicking liposomes (PM-liposomes). Both ATP and EGTA similarly reverse the *cis*-membrane interaction of synaptotagmin-1 in free [Ca^2+^] of 10 to 100 μM. High PIP_2_ concentrations in the PM-liposomes reduce the Hill coefficient of vesicle fusion and synaptotagmin-1 membrane binding; this observation suggests that local PIP_2_ concentrations control the Ca^2+^-cooperativity of synaptotagmin-1. Our data provide evidence that Ca^2+^ chelators, including EGTA and polyphosphate anions such as ATP, ADP, and AMP, electrostatically reverse the *cis*-interaction of synaptotagmin-1.

## Introduction

Exocytosis is the process of vesicle fusion and neurotransmitter release mediated by soluble *N*-ethylmaleimide-sensitive factor attachment protein receptor (SNARE) proteins, which are currently considered to be the catalysts of the fusion reaction^1, 2^. Neuronal SNARE proteins are selectively expressed in neurons and neuroendocrine cells, and regulate release of neurotransmitters and hormones^3^. Neuronal SNARE proteins consist of syntaxin-1 and SNAP-25 in the plasma membrane, and vesicle-associated membrane protein-2 (VAMP-2) (also called synaptobrevin-2) in the vesicle membrane^1^. Synaptotagmin-1 is a Ca^2+^ sensor for fast Ca^2+^-dependent exocytosis as an electrostatic switch^4^. The C2AB domain of synaptotagmin-1 coordinates Ca^2+^ binding, and the Ca^2+^-bound C2AB domain penetrates negatively-charged anionic phospholipids by electrostatic interaction^2^. Several different models of synaptotagmin-1 to describe the process of Ca^2+^-dependent vesicle fusion have been proposed, but the molecular mechanisms of synaptotagmin-1 remain controvertial^5^.

Synaptotagmin-1 is a vesicular protein and interacts with anionic phospholipids electrostatically^5^. Native vesicles contain ∼15% anionic phospholipids including phosphatidylserine (PS) and phosphatidylinositol (PI)^6^, so Ca^2+^ induces synaptotagmin-1 binding to its own vesicle membrane, i.e., the *cis-*interaction^7, 8^. Ca^2+^ fails and even slightly reduces vesicle fusion in the *in-vitro* reconstitution system, because synaptotagmin-1 preferentially interacts with vesicle membranes due to the physical proximity and this *cis*-membrane interaction prevents the *trans*-interaction of synaptotagmin-1 with the target membranes^7, 8, 9^. We have reported that ATP reverses this inactivating *cis*-interaction of synaptotagmin-1 by the electrostatic effect, and the *trans*-membrane interaction of synaptotagmin-1 only occurs to trigger vesicle fusion *in-vivo*^*10*^. This ATP effect on the *cis*-membrane interaction of synaptotagmin-1 has been confirmed independently: in a vesicle sedimentation assay a few hundred μM ATP electrostatically prevents a *cis*-configuration of synaptotagmin-1^11^, and in a fusion assay using a colloidal probe microscopy and pore-spanning membranes ATP accelerates full fusion by preventing the *cis*-interaction without affecting the trans-interaction of synaptotagmin-1^12^. However, the electrostatic regulation of the *cis*- and *trans*-membrane interaction of synaptotagmin-1 to trigger Ca^2+^-dependent vesicle fusion has not been described in detail.

Although synaptotagmin-1 is a conserved Ca^2+^ sensor for synchronous release of diverse vesicles including synaptic vesicles, large dense-core vesicles (LDCVs), and other secretory granules, the mechanism by which Ca^2+^-cooperativity is regulated is not clear. The Hill coefficient (n) in the Ca^2+^ dose-response curves for exocytosis represents Ca^2+^-cooperativity and the Hill coefficient varies depending on cell types from 2 to 5; e.g. calyx-of-Held synapses (n, 4.2)^13, 14, 15^, neuromuscular junctions (n, 3.8)^16^, bipolar cells (n, 4)^17^, pituitary melanotrophs (n, 2.5)^18^, and chromaffin cells (n, 1.8)^19^. The Hill coefficient is the intrinsic property of each cell type and factors that regulate Ca^2+^-cooperativity are poorly understood.

Synaptotagmin-1 binds to anionic phospholipids by electrostatic interaction and the Ca^2+^-binding loops of the C2 domains penetrate anionic phospholipids by reducing repulsion between anionic phospholipids and acidic residues in the C2AB domain^4^. The polybasic patch in the C2B domain electrostatically interacts with PIP_2_ in a Ca^2+^-independent manner^20^, and thereby increases the Ca^2+^-sensitivity of synaptotagmin-1 membrane binding^10, 21^. Given that the C2AB domain has five possible Ca^2+^-binding sites^22, 23^ and therefore may have the Hill coefficient up to 4∼5, but whether local PIP_2_ concentrations regulate Ca^2+^-cooperativity is not known.

Here we provide an assay to monitor the *cis*- and *trans*-membrane interaction of synaptotagmin-1 by using native LDCVs and the plasma membrane-mimicking liposomes (PM-liposomes). Ca^2+^ chelators, including EGTA and polyphosphate anions such as ATP, ADP, and AMP, electrostatically reverse the *cis*-interaction of synaptotagmin-1. Both ATP and EGTA, as Ca^2+^ chelators, have a similar effect to prevent the *cis*-membrane interaction of synaptotagmin-1 in free [Ca^2+^] of 10 to 100 μM, but ATP, which has a good buffering capacity in the range of 10 μM to 500 μM free [Ca^2+^], is an excellent Ca^2+^ buffer to study vesicle fusion and synaptotagmin-1 membrane binding. When the *trans*-membrane interaction of synaptotagmin-1 only occurs, high PIP_2_ concentrations in the PM-liposomes decrease the Hill coefficient of vesicle fusion and synaptotagmin-1 membrane binding to ∼2, suggesting that local PIP_2_ concentrations might control Ca^2+^-cooperativity of synaptotagmin-1.

## Material and Methods

### 1.1 Purification of large dense-core vesicles (LDCVs)

LDCVs, also known as chromaffin granules, were purified from bovine adrenal medullae by using continuous sucrose gradient, then resuspended in a solution of 120 mM K-glutamate, 20 mM K-acetate, and 20 mM HEPES.KOH, pH 7.4, as described elsewhere^24^.

### 1.2 Protein purification

All SNARE and the C2AB domain of synaptotagmin-1 constructs based on rat sequences were expressed in *E. coli* strain BL21 (DE3) and purified by Ni^2+^-NTA affinity chromatography followed by ion-exchange chromatography as described elsewhere^10, 20^. The stabilized Q-SNARE complex consists of syntaxin-1A (aa 183–288) and SNAP-25A (no cysteine, cysteines replaced by alanines) in a 1:1 ratio by the C-terminal VAMP-2 fragment (aa 49–96), and was purified as described earlier^25^. The C2AB domain of synaptotagmin-1 (aa 97-421) and soluble form of VAMP-2 lacking the transmembrane domain (VAMP-2_1-96_) were purified using a Mono S column (GE Healthcare, Piscataway, NJ) as described previously^26^. The stabilized Q-SNARE complex was purified by Ni^2+^-NTA affinity chromatography followed by ion-exchange chromatography on a Mono Q column (GE Healthcare, Piscataway, NJ) in the presence of 50 mM n-octyl-β-D-glucoside (OG)^10^. The point mutated C2AB domain (S342C) was labelled with Alexa Fluor 488 C5 maleimide (C2AB^A488^)^26^.

### 1.3 Lipid composition of liposomes

All lipids were obtained from Avanti Polar lipids (Alabaster, AL). Lipid composition (mol, %) of the PM-liposomes that contain the Q-SNARE complex was 45% PC (L-α-phosphatidylcholine, Cat. 840055), 15% PE (L-α-phosphatidylethanolamine, Cat. 840026), 10% PS (L-α-phosphatidylserine, Cat. 840032), 25% Chol (cholesterol, Cat. 700000), 4% PI (L-α-phosphatidylinositol, Cat. 840042), and 1% PI(4,5)P_2_ (PIP_2_, Cat. 840046). When PIP_2_ concentrations were changed, PI contents were adjusted accordingly. For FRET-based lipid-mixing assays, 1.5% 1,2-dioleoyl-sn-glycero-3-phosphoethanolamine-N-(7-nitrobenz-2-oxa-1,3-diazol-4-yl (NBD-DOPE) as a donor dye and 1.5% 1,2-dioleoyl-sn-glycero-3-phosphoethanolamine-N-lissamine rhodamine B sulfonyl ammonium salt (Rhodamine-DOPE) as an acceptor dye were incorporated in the PM-liposomes (accordingly 12% unlabelled PE).

### 1.4 Preparation of proteoliposomes

Incorporation of the Q-SNARE complex into large unilamellar vesicles (LUVs) was achieved by OG-mediated reconstitution, called the direct method, i.e. incorporation of proteins into preformed liposomes^10, 20^. Briefly, lipids dissolved in a 2:1 chloroform-methanol solvent were mixed according to lipid composition. The solvent was removed using a rotary evaporator to generate lipid film on a glass flask, then lipids were resuspended in 1.5 mL diethyl ether and 0.5 mL buffer containing 150 mM KCl and 20 mM HEPES/KOH pH 7.4. The suspension was sonicated on ice (3 × 45 s), then multilamellar vesicles were prepared by reverse-phase evaporation using a rotary evaporator as diethyl ether was removed. Multilamellar vesicles (0.5 mL) were extruded using polycarbonate membranes of pore size 100 nm (Avanti Polar lipids) to give uniformly-sized LUVs. After the preformed LUVs had been prepared, SNARE proteins were incorporated into them using OG, a mild non-ionic detergent, then the OG was removed by dialysis overnight in 1 L of buffer containing 150 mM KCl and 20 mM HEPES/KOH pH 7.4 together with 2 g SM-2 adsorbent beads. Proteoliposomes had protein-to-lipid molar ratio of 1:500.

### 1.5 Vesicle fusion assay

A FRET-based lipid-mixing assay was applied to monitor vesicle fusion *in vitro*^10, 20^. LDCV fusion reactions were performed at 37°C in 1 mL fusion buffer containing 120 mM K-glutamate, 20 mM K-acetate, 20 mM HEPES-KOH (pH 7.4), 1 mM MgCl_2_, and 3 mM ATP (**Fig.4b**). Fusion buffer in **Fig.3a,b** contains no ATP, but EGTA; 120 mM K-glutamate, 20 mM K-acetate, and 20 mM HEPES-KOH (pH 7.4), 5 mM MgCl_2_, and 10 μM EGTA. ATP should be made freshly before all experiments, because it is easily destroyed by freezing and thawing. Free Ca^2+^ concentration in the presence of Mg^2+^ and ATP or EGTA was calibrated using the MaxChelator simulation program.

**Figure 1.**
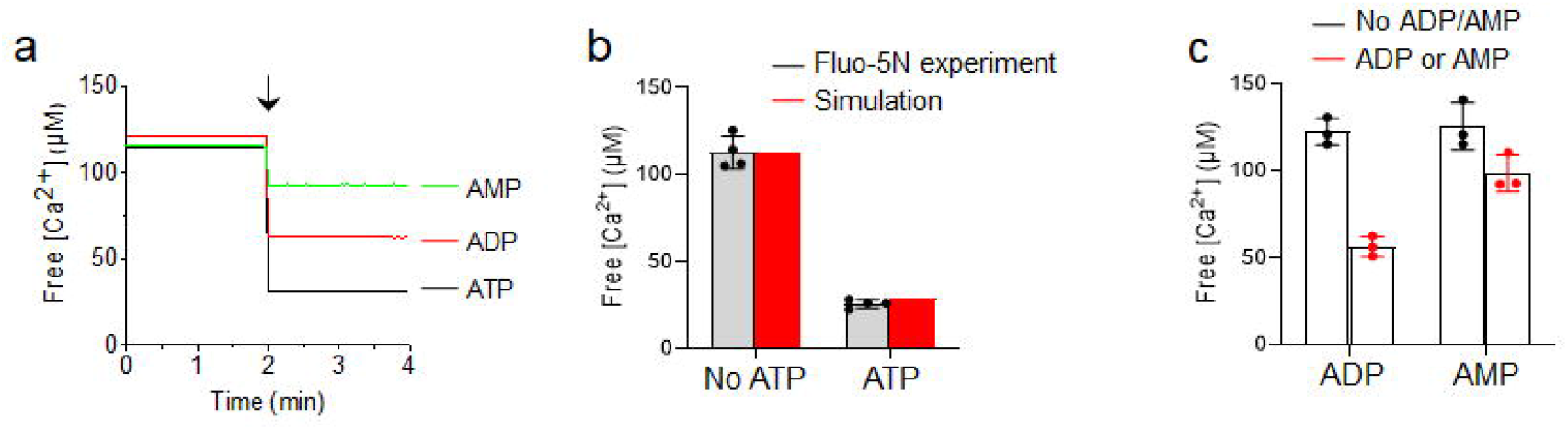
Calibration of free [Ca^2+^] using Fluo-5N and simulation in the presence of ATP. (**a**) Free [Ca^2+^] calibration in the presence of ATP, ADP, or AMP using Fluo-5N, a Ca^2+^ indicator with K_d_ = 90 μM. 5 mM of ATP, ADP, or AMP was applied (arrow). Representative trace of free [Ca^2+^] from four independent experiments. (**b**) Comparison of free [Ca^2+^] in the presence of 5 mM ATP between Fluo-5N and the MaxChelator simulation program, which calculates free [Ca^2+^] in the presence of ATP and Mg^2+^. (**c**) ADP and AMP chelate free [Ca^2+^], but the Ca^2+^-chelating efficiency is less than that of ATP. Data in **b,c** are mean ± SD from three to four independent experiments (n=3∼4).

**Figure 2.**
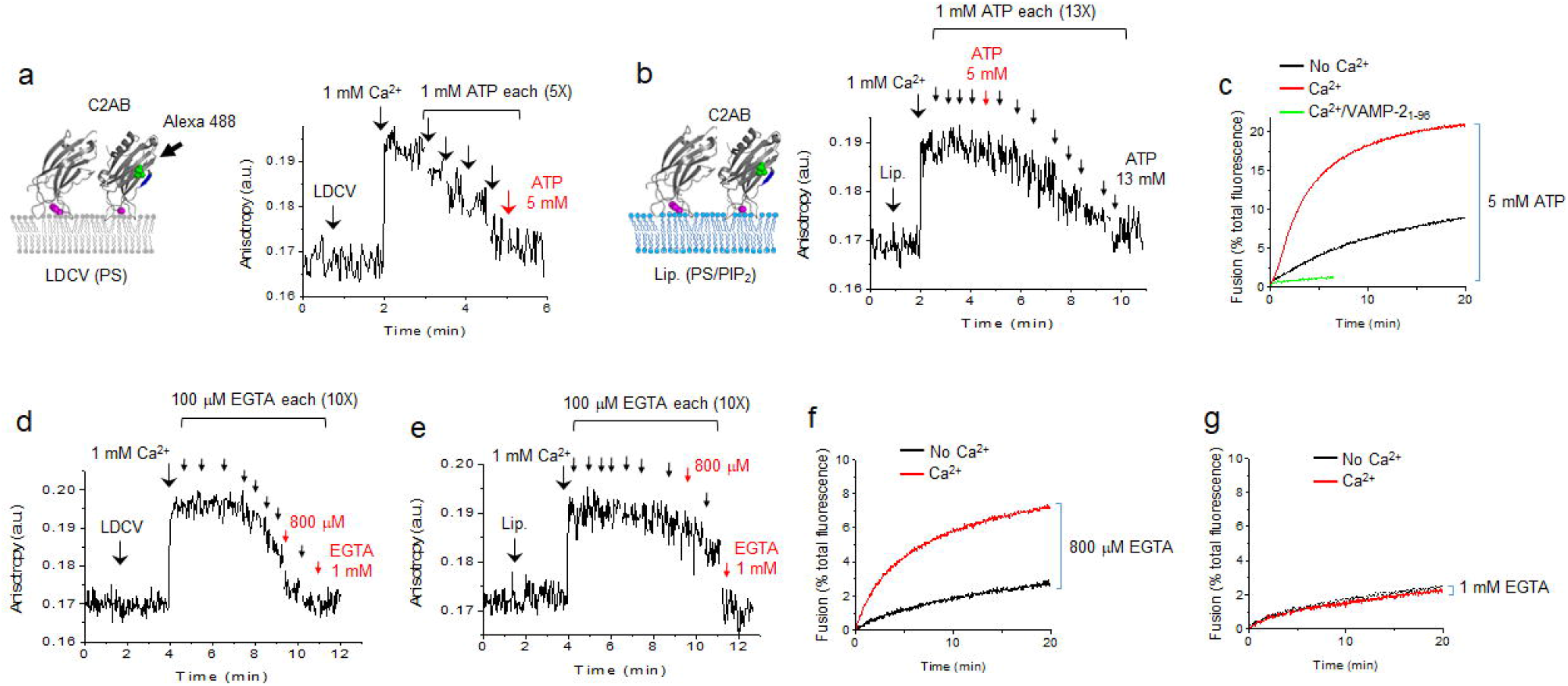
Monitoring *cis-* and *trans*-membrane interaction of synaptotagmin-1. (**a**) Binding of the C2AB domain of synaptotagmin-1 to the membrane of native vesicles (i.e., LDCVs) was monitored using fluorescence anisotropy in which the C2AB domain (Syt-1_97-421_) was labelled with Alexa Fluor 488 at S342C (green dots). (Left) The C2AB domain has Ca^2+^-binding sites (magenta) and the Ca^2+^-bound C2AB domain is inserted to membrane, thus decreasing the rotational mobility. LDCVs contain anionic phospholipids, equivalent to around 15% PS^10^. (Right) A dose of 1 mM Ca^2+^ was applied to induce binding of the C2AB domain to LDCVs, then 1 mM ATP was added five times (arrows) to reverse this binding. The final total of 5 mM ATP disrupted membrane binding of the C2AB domain (red). (**b**) C2AB domain binding to the PM-liposomes. 10% PS and 1% PIP_2_ in the PM-liposomes. 1 mM ATP was added thirteen times (arrows) and the final total of 13 mM ATP reversed membrane binding of the C2AB domain; the C2AB binding remained in 5 mM ATP (red). (**c**) *In-vitro* reconstitution of LDCV fusion using a lipid-mixing assay. Purified LDCVs were incubated with liposomes that incorporate the stabilized Q-SNARE complex. 1 mM Ca^2+^ in the presence of 5 mM ATP accelerated LDCV fusion. (**d,e**) Binding of the C2AB domain to LDCVs (**d**) and the PM-liposomes (**e**) was monitored using fluorescence anisotropy as in **a** and **b**. First, 1 mM Ca^2+^ was applied to induce binding of the C2AB domain, then 100 μM EGTA was added ten times (arrows) to reverse this binding. A total dose of 800 μM EGTA disrupted C2AB binding to LDCV (red, **d**) and a final total dose of 1 mM EGTA reversed C2AB binding to liposomes (red, **e**). (**f,g**) LDCV fusion was increased by 1 mM Ca^2+^ in the presence of 800 μM EGTA (**f**), but was not affected in the presence of 1 mM EGTA (**g**).

**Figure 3.**
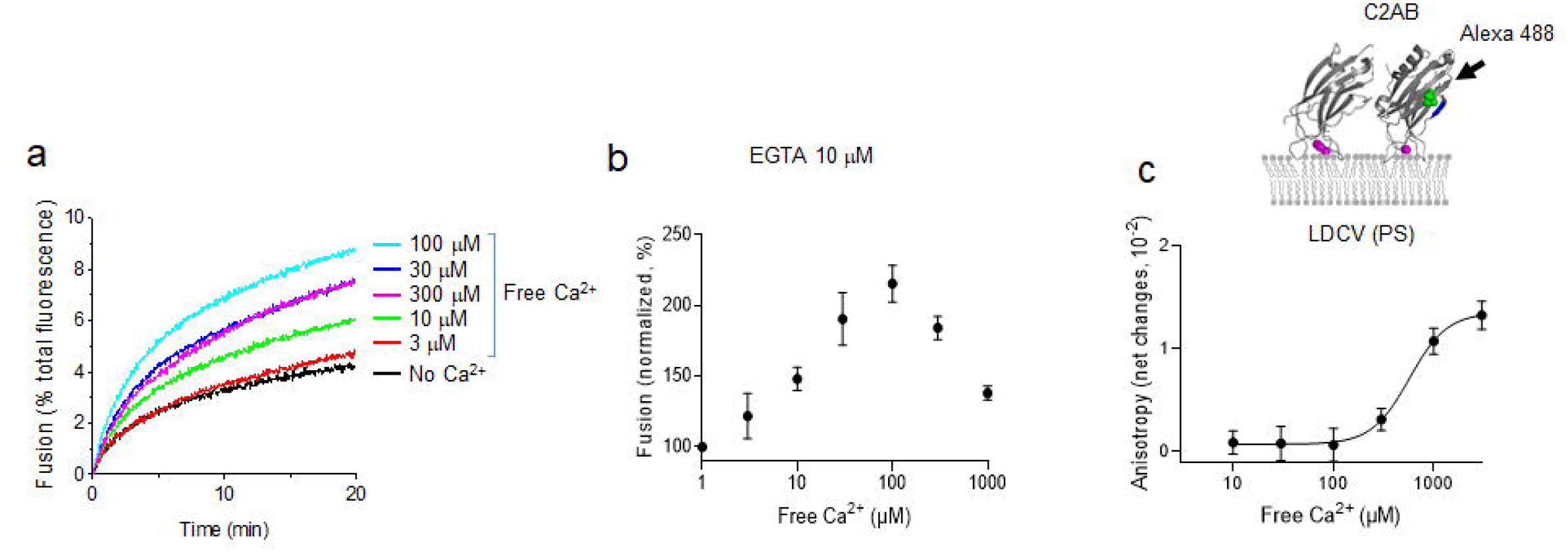
EGTA reproduces ATP effect on Ca^2+^-dependent LDCV fusion and the C2AB binding to LDCVs. (**a,b**) LDCV fusion using a lipid-mixing assay as described in **Fig. 2c** at different concentrations of Ca^2+^ in the presence of 10 μM EGTA, instead of ATP. (**a**) Representative trace of dequenching of donor fluorescence (NBD). (**b**) Dose-response curve of LDCV fusion at various free [Ca^2+^]. Fusion is normalized as a percentage of control (No Ca^2+^). (**c**) Ca^2+^ dose-response curve for C2AB binding to LDCVs in the presence of 10 μM EGTA using anisotropy as described in **Fig. 2a**. Data in **b,c** are mean ± SD from three independent experiments (n=3). Free [Ca^2+^] were calibrated using the MaxChelator simulation program. (**a-c**) ATP was not included in buffer: 120 mM K-glutamate, 20 mM K-acetate, 20 mM HEPES-KOH (pH 7.4), 5 mM MgCl_2_, and 10 μM EGTA.

**Figure 4.**
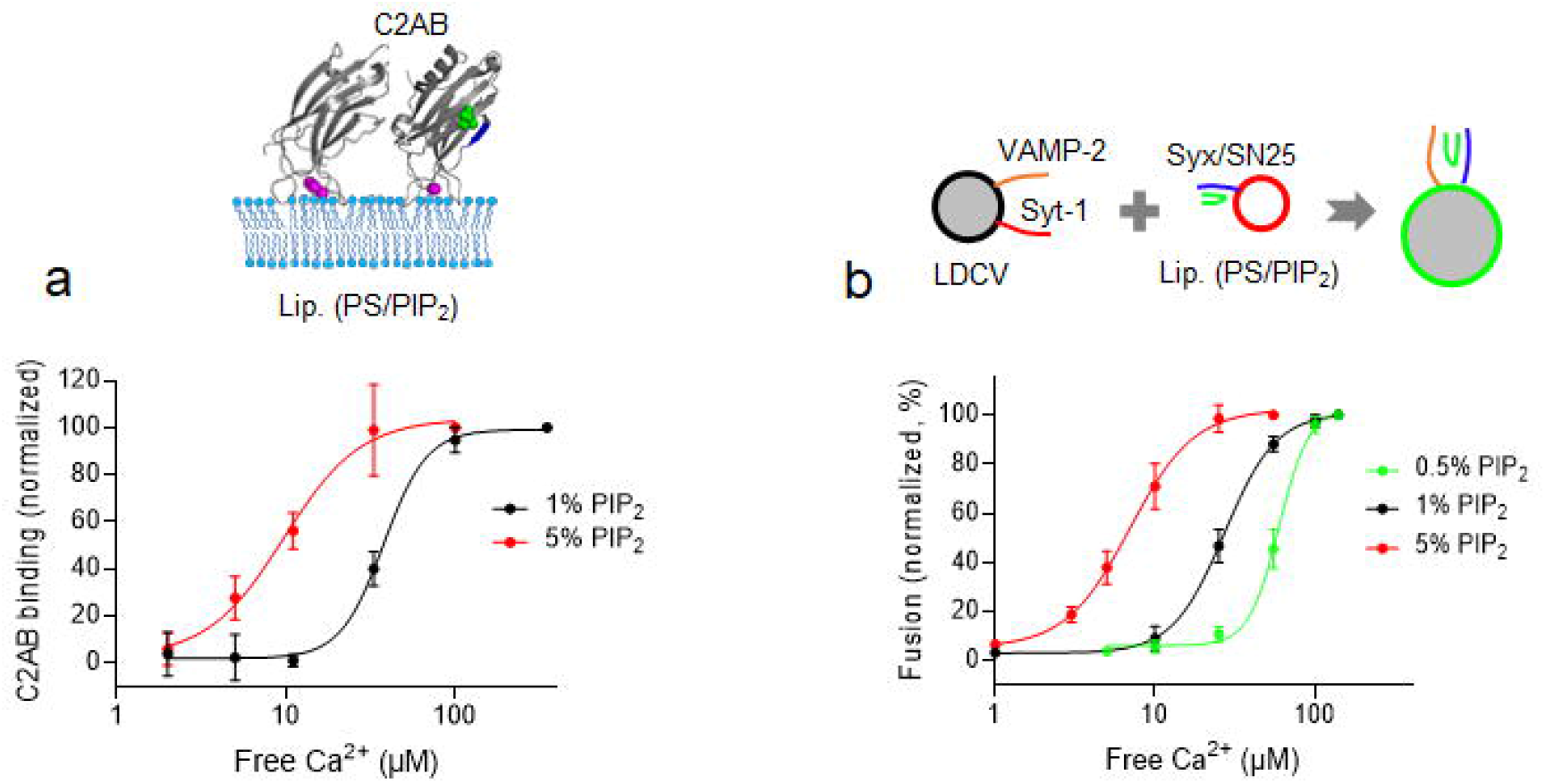
PIP_2_ concentration regulates Ca^2+^ sensitivity and cooperativity of synaptotagmin-1. (**a**) Membrane binding of the C2AB domain of synaptotagmin-1 was monitored using anisotropy as in **Fig.2b**. Ca^2+^ dose-response curve for C2AB binding to the PM-liposomes that include PS and PIP_2_. C2AB binding is presented as a percentage of maximum C2AB binding. (**b**) Ca^2+^ dose-response curve for LDCV fusion with the PM-liposomes containing different PIP_2_ concentrations. Fusion is normalized as a percentage of maximum fusion. Data in **a,b** are mean ± SD from three independent experiments (n=3). 3 mM MgCl_2_ and 1 mM ATP were included in buffer, and free [Ca^2+^] was calibrated using the MaxChelator simulation program.

The PM-liposomes that contain NBD-DOPE and Rhodamine-DOPE as a donor and an acceptor dye, respectively, were incubated with LDCVs, thus leading to dequenching of donor fluorescence (NBD) as a result of lipid dilution with unlabelled vesicle membrane^10, 20^. The fluorescence dequenching signal of vesicle fusion was measured using wavelength of 460 nm for excitation and 538 nm for emission. Fluorescence values were normalized as a percentage of maximum donor fluorescence (i.e., total fluorescence) after addition of 0.1% Triton X-100 at the end of experiments.

### 1.6 Fluorescence anisotropy measurements

The C2AB fragments (20 nM, S342C) were labelled with Alexa Fluor 488^26^. Anisotropy was measured at 37°C in 1 ml of buffer containing 120 mM K-glutamate, 20 mM K-acetate, and 20 mM HEPES-KOH (pH 7.4), 5 mM MgCl_2_, and 10 μM EGTA. First, 1 mM Ca^2+^ was applied, then ATP or EGTA was accordingly added to chelate Ca^2+^ and reverse the membrane binding of the C2AB domain; each time ATP or EGTA was uniformly mixed by pipetting and a magnetic stirring setup with dilution factor of 1:500 in 1 ml buffer. (**Fig.2**). Excitation wavelength was 495 nm and emission was measured at 520 nm. Anisotropy (*r*) was calculated using the formula *r* = (I_VV_ − G × I_VH_)/(I_VV_ + 2 ×G × I_VH_), where I_VV_ indicates the fluorescence intensity with vertically polarized excitation and vertical polarization on the detected emission and I_VH_ denotes the fluorescence intensity when using a vertical polarizer on the excitation and horizontal polarizer on the emission. G is a grating factor used as a correction for the instrument’s differential transmission of the two orthogonal vector orientations. Lipid composition of the PM-liposomes (protein-free) was identical to those used in a fusion assay except labelled PE (45% PC, 15% PE, 10% PS, 25% Chol, 4% PI, and 1% PIP_2_).

### 1.7 Ca^2+^ calibration

ATP contains negatively charged oxygen atoms which bind to Mg^2+^, Ca^2+^, or Sr^2+^, thereby chelating divalent cations^27^. Ca^2+^ concentrations were calibrated with Fluo-5N, pentapotassium salt, cell impermeant, a low-affinity Ca^2+^ indicator with a K_d_ of 90 μM. Fluo-5N (500 nM) was included in buffer containing 120 mM K-glutamate, 20 mM K-acetate, 20 mM HEPES-KOH (pH 7.4), 5 mM MgCl_2_, and 10 μM EGTA. 5 mM ATP, ADP, or AMP (sodium salt, Sigma-Aldrich) was added to chelate free Ca^2+^. The fluorescence signal was measured at 37°C with wavelength of 494 nm for excitation and 516 nm for emission. The following equation was used to measure free Ca^2+^ concentrations:

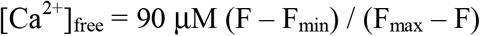

where F_min_ is the fluorescence intensity in the absence of calcium with 10 mM EGTA, F_max_ is the maxium fluorescence with 5 mM CaCl_2_, and F is the fluorescence of intermediate Fluo-5N. Fluo-5N experimental data with 5 mM ATP were correlated with the MaxChelator simulation program that calculates the free [Ca^2+^].

### 1.8 Statistical analysis

All quantitative data are mean ± SD from ≥ 3 independent experiments. Dose-response curves were fitted using four-parameter logistic equations (4PL) (GraphPad Prism) to calculate Hill slope and EC_50_.

## Results

### Calibration of free [Ca^2+^] using Fluo-5N and simulation program in the presence of ATP

Ca^2+^ is a triggering factor of vesicle fusion and intracellular Ca^2+^ concentration ([Ca^2+^]_i_) is typically ∼100 nM, but local [Ca^2+^]_i_ and Ca^2+^ microdomains at the vesicle-release sites close to voltage-gated calcium channels increase to ∼300 μM^15, 28, 29^. We used ATP, which is a low affinity Ca^2+^ buffer, to maintain ∼10 ≤ free [Ca^2+^] ≤ ∼300 μM for *in-vitro* assays^10, 20, 24, 30, 31^. ATP has a dissociation constant (K_d_) ∼230 μM [Ca^2+^]^27^, so ATP is an excellent Ca^2+^ buffer in the range of 10 μM to 500 μM free [Ca^2+^]^32, 33^. We used Fluo-5N to measure free [Ca^2+^] in the presence of ATP to confirm the predictions of [Ca^2+^] and to determine how much total [Ca^2+^] is required to achieve a desired free [Ca^2+^](**Fig.1a-c**). Fluo-5N is a low-affinity Ca^2+^ indicator (K_d_ of 90 μM)^34^, which is good for measuring around 100 μM free [Ca^2+^], because K_d_ of Ca^2+^ chelators should be close to the desired free [Ca^2+^]^35^. EGTA (10 μM) was included to remove contaminating Ca^2+^ for the calibration of free [Ca^2+^]. An initial total 113 μM free [Ca^2+^] was reduced to 26 μM in the presence of 5 mM ATP by its chelation of Ca^2+^ (**Fig.1a,b**).

Then we compared this experimental data of free [Ca^2+^] with the MaxChelator, which is a computer simulation programs^32, 35^ that enables calculation of appropriate stoichiometric concentrations of Ca^2+^ and Mg^2+^ in the presence of different Ca^2+^ chelators such as EGTA and ATP, and thereby provides detailed infomation to obtain the desired free [Ca^2+^]^35^. The MaxChelator program included 5 mM Mg^2+^ and 10 μM EGTA, and assumed 37°C as in the Ca^2+^ calibration experiments (**Fig.1a**). Indeed the MaxChelator calculated free [Ca^2+^] = 29 μM in the presence of 5 mM ATP with 113 μM total [Ca^2+^] at pH 7.4. This agreement with the measured free [Ca^2+^] = 26 μM confirms that the MaxChelator can predict free [Ca^2+^] obtained in experiments that use a Fluo-5N fluorescent Ca^2+^ indicator (**Fig.1b**).

Negatively-charged oxygen atoms of ATP chelate divalent cations such as Mg^2+^, Ca^2+^, or Sr^2+27^. In the experiments, 5 mM ADP or 5 mM AMP chelated Ca^2+^, thereby reducing free [Ca^2+^] from 122 μM to 57 μM and from 126 μM to 99 μM, respectively (**Fig.1c**). Increasing the number of phosphate groups in Adenosine increases Ca^2+^ affinity and lowers K_d_ by increasing the number of Ca^2+^ ions that are bound^27, 33^. ATP, ADP, and AMP have distinct ranges of Ca^2+^-buffering capacity and distinct K_d_ values^33^, so Ca^2+^-chelating effect is ATP > ADP > AMP (**Fig.1a-c**). Altogether, the predictions of free [Ca^2+^] in the complex buffer solutions including Mg^2+^, ATP and EGTA were confirmed using a fluorescent Ca^2+^ indicator.

### Monitoring the *cis*- and *trans*-membrane interaction of synaptotagmin-1

Synaptotagmin-1 interacts with anionic phospholipids by electrostatic interaction. Native vesicles contain ∼15% anionic phospholipids, including phosphatidylserine (PS) and phosphatidylinositol (PI)^6^. Therefore, Ca^2+^ induces synaptotagmin-1 to bind to its own vesicle membrane, i.e., *cis-*interaction, which prevents *trans*-interaction to the plasma membranes and thereby inactivates the ability of synaptotagmin-1 to trigger fusion^7, 8, 9^.

Ca^2+^-bound synaptotagmin-1 is inserted to native vesicle membranes such as synaptic vesicles and large dense-core vesicles (LDCVs) that contain anionic phospholipids ^10^. However, ATP electrostatically prevents the *cis-*interaction of synaptotagmin-1, whereas the *trans*-interaction of synaptotagmin-1 to the plasma membrane remains active to mediate Ca^2+^-dependent vesicle fusion, because PIP_2_ overcomes the inhibitory effect of ATP by increasing the membrane-binding affinity of the C2AB domain ^10, 11, 12^.

We tested an assay that uses fluorescence anisotropy measurement to monitor the *cis*- and *trans*-membrane interaction of synaptotagmin-1 (**Fig.2**). Direct measurement of the *cis*- and *trans*-membrane interaction of endogenous synaptotagmin-1 in native vesicle membranes is impossible, so we monitored the binding of an exogenously-added C2AB domain of synaptotagmin-1 (Syt_97-421_), which was labelled with Alexa Fluor 488 at S342C (**Fig.2a**). We took advantage of a single fluorescent labelling for anisotropy measurement to monitor the interaction of the C2AB domain with native vesicles or liposomes; the membrane-bound C2AB domain leads to increase of fluorescence anisotropy due to a reduction in the rotational mobility^10^ (**Fig.2a,b**). It is noted that our experiments using the cytoplasmic C2AB domain are intended to shed light on the *cis*- and *trans*-interactions, but the geometry is not truly being imitated.

We first monitored the *cis*-membrane interaction between the C2AB domain and the LDCV membranes (**Fig.2a**). The presence of 1 mM Ca^2+^ increased fluorescence anisotropy; this change indicates the C2AB domains bind to LDCV membranes in a Ca^2+^-dependent manner. Five sequential applications of 1 mM ATP gradually decreased the anisotropy signal by chelating Ca^2+^; this result suggests dissociation of the C2AB domain from LDCVs (**Fig.2a**). 5 mM ATP in the presence of 1 mM Ca^2+^ almost completely disrupted the *cis*-membrane interaction of the C2AB domain with the LDCV membranes (**Fig.2a**); free [Ca^2+^] in the presence of Mg^2+^, ATP and EGTA was calibrated using the MaxChelator simulation program and free [Ca^2+^] was 351 µM in case of 5 mM ATP and 1 mM Ca^2+^ (**Table 1**).

**Table 1.** Calibration of free Ca^2+^ concentration in the presence of ATP or EGTA^a^.

| ATP concentration<br>[mM] | Free $\text{Ca}^{2+}$ [$\mu\text{M}$]<br>(total 1 mM $\text{Ca}^{2+}$)<sup>b</sup> | EGTA concentration<br>[$\mu\text{M}$] | Free $\text{Ca}^{2+}$ [$\mu\text{M}$]<br>(total 1 mM $\text{Ca}^{2+}$) |
| --- | --- | --- | --- |
| 1 | 896 | 100 | 900 |
| 2 | 785 | 200 | 800 |
| 4 | 508 | 300 | 700 |
| 5 | 351 | 800 | 200 |
| 13 | 31 | 1000 | 12 |

<sup>a</sup>, The MaxChelator simulation program was used to calculate free $\text{Ca}^{2+}$ concentration in the presence of ATP or EGTA.
<sup>b</sup>, 1 mM $\text{Ca}^{2+}$ , 5 mM $\text{Mg}^{2+}$ , and 10 $\mu\text{M}$ EGTA.

Next, we tested the *trans*-membrane interactions between the C2AB domain and the PM-liposomes; 10% PS, 4% PI, and 1% PIP_2_ were included in the PM-liposomes (**Fig.2b**). The C2AB domain of synaptotagmin-1 bound to liposomes in response to 1 mM Ca^2+^, and this *trans*-membrane interaction was reduced by ATP, 1 mM applied thirteen times sequentially (**Fig.2b**). Free [Ca^2+^] in different ATP concentrations was summarized in **Table 1**. Ca^2+^-dependent vesicle fusion is accelerated by the increase of the *trans*-interactions and the decrease of the *cis*-membrane interaction of synaptotagmin-1^10, 20^, so we hypothesized that 5 mM ATP in the presence of 1 mM Ca^2+^ is appropriate to observe Ca^2+^-dependent fusion (red in **Fig.1a,b**). To test this hypothesis and examine the effect of the *cis*- and *trans*-membrane interaction of synaptotagmin-1 on vesicle fusion, we applied a reconstitution system of vesicle fusion by using native LDCVs^10, 20, 24^. The PM-liposomes contain the stabilized Q-SNARE complex (syntaxin-1A and SNAP-25A in a 1:1 molar ratio^25^).

Indeed, 5 mM ATP in the presence of 1 mM Ca^2+^ (i.e., 351 µM free [Ca^2+^] according to the MaxChelator program (**Table 1**)) dramatically accelerated LDCV fusion, which was completely blocked by the soluble VAMP-2 (VAMP-2_1-96_); this results indicates SNARE-dependent vesicle fusion (**Fig.2c**). We have previously shown that 300∼400 μM free [Ca^2+^] in the absence of ATP fails to enhance vesicle fusion, but rather slightly inhibits fusion, because the *cis*-membrane interaction of the C2AB domain to native vesicle membranes becomes robust from 100 μM up to 3 mM^10^. ATP prevents this *cis*-membrane interaction by charge screening and competing with the vesicle membrane, thus allowing synaptotagmin-1 to interact in *trans* with the plasma membrane^10^.

Polyphosphates such as ATP reverse an inactivating *cis*-interaction of synaptotagmin-1 by an electrostatic effect (**Fig.2a-c**). Next, we tested whether other Ca^2+^ chelators, e.g., EGTA, can have a similar inhibitory effect on the *cis*-membrane interaction. Anisotropy measurement was performed to monitor the *cis*- and *trans*-membrane interaction of the C2AB domain (**Fig.2a,b**). EGTA was applied 10 times (100 μM each in the presence of 1 mM Ca^2+^) to reverse the *cis*-interaction of the C2AB domain to LDCVs (**Fig.2d**). Application of 800 μM EGTA dramatically disrupted the *cis*-interaction in the presence of total 1 mM Ca^2+^ (red in **Fig.2d**); free [Ca^2+^] was 200 μM (**Table 1**). However, the *trans*-membrane interactions of the C2AB domain to the PM-liposomes remained robust in the presence of 800 μM EGTA with 1 mM Ca^2+^ (200 μM free [Ca^2+^], **Fig.2e**), whereas 1 mM EGTA significantly disrupted both the *cis*- and *trans*-membrane interactions of the C2AB domain (**Fig.2d,e**); free [Ca^2+^] was 12 μM (**Table 1**).

Anisotropy measurement is useful to find a Ca^2+^-buffering condition to observe Ca^2+^-dependent vesicle fusion, where the *cis*-membrane interaction is prevented and the *trans*-interaction remains active. The presence of 800 μM EGTA with 1 mM Ca^2+^ (200 μM free [Ca^2+^], **Table 1**) significantly reversed the *cis*-interaction (**Fig.2d**), but had a minor effect on the *trans*-interaction (**Fig.2e**). Indeed, 800 μM EGTA with 1 mM Ca^2+^ reproduced Ca^2+^-dependent LDCV fusion (**Fig.2f**). 1 mM EGTA with 1 mM Ca^2+^ (12 μM free [Ca^2+^], **Table 1**) failed to accelerate fusion, because the *trans*-interaction of the C2AB domain was dramatically disrupted by 1 mM EGTA (red in **Fig.2e**); it is mainly because of low free [Ca^2+^]. Taken together, we established an anisotropy assay to monitor the *cis*- and *trans*-membrane interaction of synaptotagmin-1 by using native LDCVs and the PM-liposomes. Our data suggest that Ca^2+^ chelators such as EGTA, in addition to polyphosphates such as ATP, can prevent the *cis*-membrane interaction of synaptotagmin-1 by the electrostatic effect in a certain range of free [Ca^2+^].

### EGTA reproduces the biphasic regulation of Ca^2+^ on LDCV fusion

We have previously reported the biphasic regulation of Ca^2+^ on LDCV fusion; 10-100 μM free Ca^2+^ exponentially accelerates native vesicle fusion, but > 300 μM free [Ca^2+^] progressively reduces Ca^2+^-dependent fusion, showing biphasic regulation of Ca^2+^ on LDCV fusion in a bell-shaped dose-dependence^20^. ATP was used for Ca^2+^-buffering to maintain free [Ca^2+^] in the range of 10 μM to 500 μM^20^. We examined whether EGTA reproduces the biphasic regulation of Ca^2+^ on LDCV fusion (**Fig.3a,b**). Instead of ATP, 10 μM EGTA was included in fusion buffer and free [Ca^2+^] was calculated using the MaxChelator program. As expected, biphasic regulation of Ca^2+^ on LDCV fusion was observed, where Ca^2+^-dependent fusion progressively increased until [Ca^2+^] = ∼100 μM, and gradually decreased at [Ca^2+^] from 300 μM to 1 mM (**Fig.3a,b**).

Biphasic regulation of Ca^2+^ on LDCV fusion is mediated by two different mechanisms: i) millimolar range of [Ca^2+^] decreases the *trans*-interaction of synaptotagmin-1 by shielding PIP_2_ and ii) sub-millimolar range of [Ca^2+^] above 300 μM increases the *cis*-interaction of synaptotagmin-1 to its own vesicle membrane^20^. To further confirm the *cis*-interaction at higher [Ca^2+^], we performed anisotropy measurement (**Fig.2a,d**) to study the Ca^2+^ dose-response of the *cis*-interaction of synaptotagmin-1 in the presence of EGTA instead of ATP (**Fig.3c**). Indeed, the *cis*-membrane interaction of the C2AB domain gradually increased from 300 μM [Ca^2+^] and remained robust at millimolar [Ca^2+^](**Fig.3c**). Note that ATP and EGTA give rise to slightly different kinetics of the Ca^2+^ dose-response curves of vesicle fusion and the *cis*-interaction of synaptotagmin-1^20^, because ATP effectively buffers free [Ca^2+^] in the range of 10 μM to 500 μM, but EGTA cannot efficiently buffer free [Ca^2+^] in this range.

### PIP_2_ concentration regulates Ca^2+^ cooperativity of synaptotagmin-1

Synaptotagmin-1 binds to anionic phospholipids by electrostatic interaction and the Ca^2+^-binding loops of the C2 domains are inserted to anionic phospholipids in a Ca^2+^-dependent manner; aspartate residues of the Ca^2+^-binding loops in the C2-domains together with anionic membrane lipids coordinate Ca^2+^-ions^21, 23, 36^. PIP_2_ enhances Ca^2+^-sensitivity of synaptotagmin-1 by interacting with the polybasic patch in the C2B domain^10, 21^. Ca^2+^-cooperativity of synaptotagmin-1 varies among cell types, with the Hill coefficients ranging from ∼2 to ∼5. We tested that PIP_2_ also regulates Ca^2+^-cooperativity of synaptotagmin-1 for membrane binding (**Fig. 4a, Table 2**) and vesicle fusion (**Fig. 4b, Table 2**). Increases of PIP_2_ concentration from 1% to 5% in the PM-liposomes shifted Ca^2+^ titration curves for membrane binding to the left side; this change indicates increased Ca^2+^ sensitivity, but reduced Ca^2+^ cooperativity (**Fig. 4a, Table 2**).

**Table 2.**
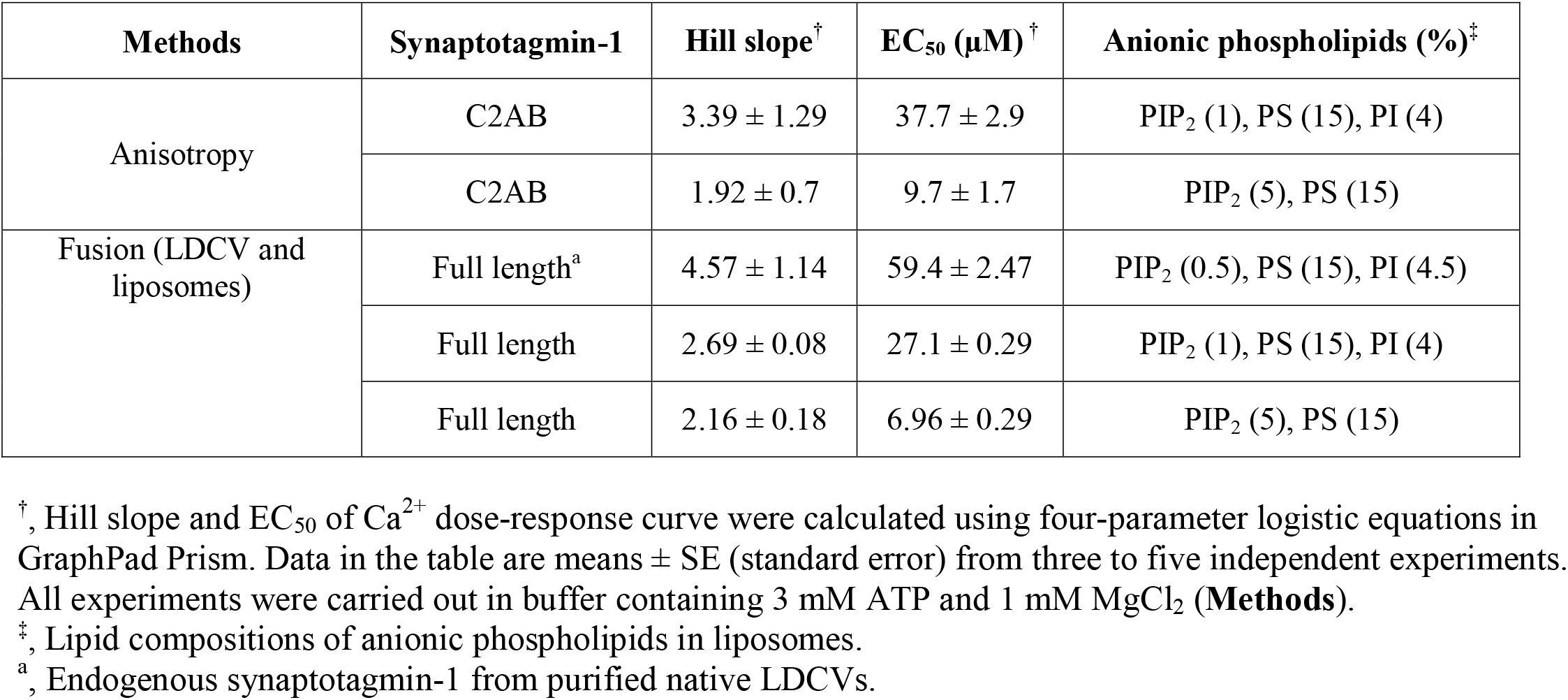
Hill slope and EC_50_ of Ca^2+^ dose-response curve.

Next, we observed that Ca^2+^-cooperativity of synaptotagmin-1 for vesicle fusion was also reduced by increasing PIP_2_ concentration, correlating with the Ca^2+^-cooperativity of synaptotagmin-1 for membrane binding. The Ca^2+^ dose-response curve for LDCV fusion was shifted leftward as PIP_2_ concentration was increased in the PM-liposomes (**Fig. 4b, Table 2**). Taken together, high PIP_2_ concentration increases the sensitivity of synaptotagmin-1 to Ca^2+^, but lowers Ca^2+^ cooperativity. These changes imply that increasing the negative electrostatic potential in the plasma membranes attracts Ca^2+^-bound synaptotagmin-1 with low Ca^2+^ cooperativity, in which the total numbers of Ca^2+^ ions coordinated to one synaptotagmin-1 might be reduced to 2 ∼ 3 (see **Discussion**).

## Discussion

The *cis*-binding of synaptotagmin-1 occurs in native vesicles such as LDCVs and synaptic vesicles, and inactivates Ca^2+^-dependent vesicle fusion by preventing the *trans*-interaction of synaptotagmin-1. Independent groups have confirmed that ATP at physiological concentrations disrupts such *cis*-interaction of synaptotagmin-1^11, 12, 37^. Here we show that Ca^2+^ chelators, including EGTA and polyphosphate anions such as ATP, ADP, and AMP, electrostatically reverse the *cis*-interaction of synaptotagmin-1. We propose that Ca^2+^ chelators compete with vesicle membranes that contain anionic phospholipids in binding to Ca^2+^ and disrupt the *cis*-interaction of synaptotagmin-1 by charge screening^10^. However, PIP_2_ overcomes this inhibitory effect of ATP, because PIP_2_ enhances the Ca^2+^-binding affinity of synaptotagmin-1^21, 38^; this high Ca^2+^ affinity of the C2AB domain to PIP_2_-containing membranes is not affected by ATP^10^.

EGTA and 1,2-bis(o-aminophenoxy)ethane-N,N,N0,N0-tetraacetic acid (BAPTA) are well-known and reliable Ca^2+^ buffers in the range of 10 nM to 1 μM [Ca^2+^] at the typical intracellular pH of 7.2^33, 35^. Given that EGTA and BAPTA have a K_d_ of 67 nM and 192 nM [Ca^2+^] at pH 7, respectively, and have a higher affinity for Ca^2+^ than for Mg^2+35^, both EGTA and BAPTA effectively buffer free [Ca^2+^] only at concentrations < 1 μM^33, 39^, which is close to intracellular free [Ca^2+^]. However, EGTA is sensitively dependent on pH,^35^ and BAPTA family has a strong dependence on ionic strength^40^; importantly, because EGTA and BAPTA have nanomolar-level *K*_d_, they poorly buffer free [Ca^2+^] in the range of 10 to 500 μM. In contrast, ATP has K_d_ 230 µM^27^ and is an excellent buffer for free [Ca^2+^] in the range of 10 to 500 μM^33^. Synaptotagmin-1 is a low-affinity Ca^2+^ sonsor; 10 to 100 μM [Ca^2+^] exponentially induce synaptotagmin-1 binding to membrane that contain PS and PIP_2_ with K_d_ ∼50 μM^21, 26^. Therefore, ATP is an appropriate and better Ca^2+^ buffer than EGTA or BAPTA to study the synaptotagmin-1 activity to bind membrane and trigger vesicle fusion. Indeed, we oberserved that ATP and EGTA result in slightly different kinetics of the Ca^2+^ dose-response curves of vesicle fusion and of the *cis*-interaction of synaptotagmin-1^10, 20^ (**Fig. 3b,c**), because ATP has a different Ca^2+^-buffering capacity than EGTA.

The K_d_ of low-affinity Ca^2+^ indicator dyes can vary depending on ionic strength and is changed by anions such as ATP^41^; e.g., the K_d_ of low-affinity Ca^2+^ indicator dyes is increased by ATP and slightly decreased by excess Mg^2+^. The K_d_ of Fluo-5N can be altered by the presence of ATP/Mg^2+^, which makes it difficult to accurately measure free [Ca^2+^]. ATP binds both Ca^2+^ and Mg^2+^ with a different affinity^27, 33^, so computer simulation programs^32, 35^ like the MaxChelator are useful to calibrate free [Ca^2+^] in the presence of Mg^2+^, ATP or EGTA by calculating free [Mg^2+^], [Ca-ATP], and [Mg-ATP]^35^. We confirmed the MaxChelator-based predictions using a Fluo-5N fluorescent Ca^2+^ indicator (**Fig.1b**).

Both the C2A and C2B domains of synaptotagmin-1 have highly cooperative Ca^2+^-dependent binding to membranes that contain anionic phospholipids^26, 42, 43, 44, 45^. Furthermore, synaptotagmin-1 contains a polybasic region within the C2B domain that binds to PIP_2_ in an Ca^2+^-independent manner^46, 47^ and enhances Ca^2+^ sensitivity of synaptotagmin-1 membrane binding^21^ and exocytosis^48^. The C2AB domain has five possible Ca^2+^-binding sites^22, 23^; negatively charged oxygen atom from acidic aspartate residues in the C2AB domain and negatively charged oxygen atom from anionic phospholipids provide complete coordination sites for Ca^2+23, 36^. Ca^2+^ cooperativity of the C2AB domain seems reasonable when the Hill coefficient is ∼4 to 5, but what regulates Ca^2+^ cooperativity remains poorly understood, e.g., low Hill coefficient (n, 2∼3) in neuroendocrine cells such as pituitary melanotrophs (n, 2.5)^18^ and chromaffin cells (n, 1.8)^19^, but high Hill coefficient in synapses including calyx-of-Held synapses (n, 4.2)^13, 14, 15^, neuromuscular junctions (n, 3.8)^16^, and bipolar cells (n, 4)^17^. We overserved that increasing PIP_2_ concentration reduces the Hill coefficient, which represents Ca^2+^ cooperativity (**Fig. 4**). Our data support that local PIP_2_ concentration might control Ca^2+^ cooperativity by allosterically-stabilized dual binding of synaptotagmin-1 to Ca^2+^ and PIP_2_^38^.

In this study, we investigate the electrostatic regulation of C2AB binding to vesicle membrane and the PM-liposomes. We have previously observed that Ca^2+^-independent interactions of the C2AB domain with the PM-liposomes containing anionic phospholipids (10% PS/1% PIP_2_) is significantly disrupted in the presence of physiological concentration of ATP/Mg^2+^, but this Ca^2+^-independent interaction remains strong when the PM-liposomes contain high PIP_2_ (10% PS/5% PIP_2_), suggesting that high PIP_2_ concentrations are required for Ca^2+^-independent binding of the C2AB domain in physiological ionic strength^20^. Here, we have used 10% PS/1% PIP_2_ in the PM-liposomes to selectively examine the Ca^2+^-dependent membrane interaction and binding of the C2AB domain. However, in the pre-fusion state for vesicle docking and priming, the C2AB domain of synaptotagmin-1 is most likely bound to the plasma membrane through the PIP_2_-interacting polybasic region of the C2B domain and the SNARE complex^49^ in a Ca^2+^-independent manner. Ca^2+^ can induce a re-orientation of the C2AB domain on the plasma membrane by changing the binding mode with the SNARE complex^49^ and PIP_2_^45^. This change in orientation may act as a switch to trigger synaptotagmin-1-dependent vesicle fusion in neurons and neuroendocrine cells. Our results do not rule out the possibility that Ca^2+^-independent interactions of synaptotagmin-1 with the SNARE complex^49^ and it remains a topic of further study to include Ca^2+^-independent interactions of synaptotagmin-1 in our system for physiological relevance.

## Acknowledgements

This work was supported by the grant from Qatar Biomedical Research Institute (Project Number SF 2019 004 to Y.P.).

